# Impact of Sex on Heroin Intravenous Self-Administration by Heterogeneous Stock Rats

**DOI:** 10.64898/2026.04.08.717349

**Authors:** Michael A. Taffe, Sydney L. Mehl, Sara R. M. U. Rahman, Yanabel Grant

## Abstract

**Background:** Intravenous self-administration (IVSA) of opioids by rats has been shown frequently to exhibit no sex differences, in many cases a higher intake of females, and only rarely higher rates in males. A diversity of methodological parameters (opioid identity, training doses, rat strain, session duration) makes it difficult to identify consistent contributions to these outcomes.

**Objective:** To determine if Heterogeneous Stock (HS) rats derived from 8 founder strains differ by sex in the IVSA of opioids.

**Methods:** Male and female Heterogeneous Stock (N=7-8 per sex) rats were permitted to self-administer heroin (20 µg/kg/infusion) in 2 hour sessions under a Fixed Ratio 1 schedule of reinforcement. After acquisition, animals completed sessions in which different infusion doses of heroin (0, 15, 30, 60, 120 µg/kg/infusion), oxycodone (0, 30, 60, 150, 300 µg/kg/infusion) and fentanyl (0, 0.625, 1.25, 2.5, 5.0 µg/kg/infusion) were assessed. Next, animals were evaluated on doses of heroin (15, 30, 60, 120 µg/kg/infusion), oxycodone (30, 60, 150, 300 µg/kg/infusion) and fentanyl (0.625, 1.25, 2.5, 5.0 µg/kg/infusion) under a Progressive Ratio schedule. Anti-nociceptive effects of heroin (0.56-2.4 mg/kg, s.c.) were examined with a warm water tail-withdrawal assay.

**Results:** Female HS rats consistently self-administered more infusions of opioids, including heroin during acquisition, all three opioids during FR-1 dose substitution and of oxycodone and fentanyl in the PR procedure. Male rats were moderately more sensitive to the anti-nociceptive effects of heroin.

**Conclusions:** Female rats drawn at random from a genetically diverse population self-administer opioids at higher rates than their male counterparts.

## Introduction

Prior to the introduction of the U.S. National Institutes of Health’s mandate (Clayton and Collins, 2014; Shansky and Woolley, 2016) to consider sex as a biological variable (SABV) the scientific literature on the intravenous self-administration (IVSA) of drugs directly compared the sexes only sporadically. Despite the use of female rats in theoriginal demonstration of IVSA (Weeks, 1962), a majority of studies have focused only on male rats. Some early reports found no sex-difference in heroin IVSA intake, albeit female rats reached training arbitrary training criteria more rapidly (Carroll et al., 2001; Lynch and Carroll, 1999). In rats selectively bred for high saccharin preference, however, females self-administered more heroin than did males (Carroll et al., 2002). More recent reports on the IVSA of heroin, oxycontin or fentanyl have sometimes reported no sex differences in group mean intake across acquisition (Carter et al., 2021; D’Ottavio et al., 2023; Lynch and Carroll, 1999; Mavrikaki et al., 2021; Mavrikaki et al., 2017; Rakowski et al., 2025), in other cases it has been shown that females will self-administer more (Cicero et al., 2003; Deckers et al., 2025; Klein et al., 1997; Nguyen et al., 2020; Taffe et al., 2026). This makes it clearly premature to suggest that a single outcome of no-difference makes it unnecessary to power studies to determine sex differences (D’Ottavio et al., 2023).

It can be difficult to determine the experimental variables which may be critical to the presence of sex differences given variation in opioid drug, session durations, acquisition intervals, rat strain, light cycle, training dose and post-acquisition experimental manipulations. For example, one report showed that sex differences were observed at lower training doses of fentanyl but not at higher doses (Towers et al., 2022), which is similar to a recent report that changing the per-infusion dose of methamphetamine to 0.025 mg/kg introduced a sex difference that was not observed at 0.05 mg/kg (Gutierrez et al., 2025). Rat strain may be critical, indeed studies have reported strain differences in IVSA of heroin by female (Schmidt et al., 2021) and male (Lecca et al., 2020) rats that equal or exceed the magnitude of sex differences reported within strain.

Previous reports tend to use one of the strain of rats typical of laboratory studies, with a particular dominance of Sprague-Dawley and Wistar rats. (Note that nominal Sprague-Dawley rats from different sources may differ from each other on behavioral phenotypes (Fitzpatrick et al., 2013), thus it is advantageous to consider different source Sprague-Dawley as potentially different strains.) Heterogeneous Strain (HS) rats were created (Hansen and Spuhler, 1984) to provide a genetically diverse population suitable for genome-wide association studies. The HS rats were derived from interbreeding eight inbred rat strains (ACI/N, BN/SsN, BUF/N, F344/N, M520/N, MR/N, WKY/N, and WN/N) and are maintained with breeding strategy designed to minimize inbreeding and genetic drift (Solberg Woods and Palmer, 2019). Studies using traditional group sizes of these HS rats may therefore offer an opportunity to determine when behavioral effects are likely to generalize beyond traditional rat strains, given the genetic diversity represented in any random sample from the population.

Most of the data for heroin IVSA by HS rats has been generated using a 20 ug/kg/infusion training dose (Kuhn et al., 2025a; Rakowski et al., 2025). The genome-wide association study was intended to identify genetic signatures with relevance to heroin use disorders based on IVSA data from ∼1,000 rats, thus a standardized methodological approach was employed. No sex difference were reported across fifteen 12 h Long Access (LgA) acquisition sessions (Greenberg et al., 2024) This is consistent with a recent report using a small sample of HS rats in which no sex differences were observed in ten acquisition 2 h Short Access (ShA) sessions (Rakowski et al., 2025). This latter found, however, that ***males*** self-administered more heroin in a subsequent 15 sessions where ∼continuous drug access was increased to 4 hours. Prior work includes training heroin IVSA with 20 µg/kg/infusion (Ahmed et al., 2000; Kenny et al., 2006), 30 µg/kg/infusion (Hyytia et al., 1996), 60 µg/kg/infusion (Carrera et al., 1999; Ettenberg et al., 1982; Vendruscolo et al., 2011) and 75 µg/kg/infusion (D’Ottavio et al., 2023) training doses of heroin, thus the training dose (20 ug/kg/infusion) used in the HS rat studies is on the lower end. Dose may not always be important, indeed, a direct comparison of male rats trained on 30, 60 and 120 µg/kg/infusion found very little difference in the number of infusions earned during 12 hour sessions (Wade et al., 2015). Nevertheless it is possible the identification of sex differences depends on training dose.

The goal of the present investigation was to determine if young adult, HS rats exhibit a sex difference in the self-administration of heroin at a training dose (20 ug/kg/infusion) that is on the lower end for heroin IVSA studies. Our prior work showed female Wistar young adult rats self-administer more infusions of heroin (60 µg/kg/infusion) in 2 h sessions (Taffe et al., 2026) and more oxycodone (150 µg/kg/infusion) in 8 h sessions (Nguyen et al., 2020) than do their male counterparts, thus our hypothesis is that HS females will IVSA at higher rates. Experiments compared the sexes on heroin IVSA acquisition, on the impact of dose substitution of heroin under Fixed Ratio 1 and Progressive Ratio schedules of reinforcement, on the impact of dose substitution of oxycodone and fentanyl and on sensitivity to pre-treatment with the opioid antagonist naloxone. Additional experiments compared the sexes on the anti-nociceptive effects of non-contingent heroin to further determine any potential sex differences in opioid sensitivity.

## Methods

### Subjects

Male and Female Heterogeneous Stock (N=8 per sex) bred at the University of California, San Diego by the Dr. Abraham A. Palmer laboratory (McwiWfsmAap:HS #155269102, RRID:RGD_155269102) were transferred to the Taffe laboratory at 9-11 weeks of age. HS rats were derived from interbreeding eight inbred rat strains (ACI/N, BN/SsN, BUF/N, F344/N, M520/N, MR/N, WKY/N, and WN/N) and maintained with a large number of breeder pairs using a randomized breeding strategy, with each pair contributing one male and one female per generation to minimize inbreeding and genetic drift (Solberg Woods and Palmer, 2019). Animals were provided from one breeding cohort and were not selected based on genotype. Individual dates of birth stretched across a twelve day interval, thus the center of weeks of age are described for the experimental procedures, see below. The vivarium was kept on a 12:12 hour reversed light-dark cycle (lights out at 0800); studies were conducted during vivarium dark. Food and water were provided ad libitum in the home cage. Procedures were conducted in accordance with protocols approved by the IACUC of the University of California, San Diego and were consistent with recommendations in the NIH Guide (Garber et al., 2011).

### Drugs

Heroin (Diamorphine HCl), oxycodone HCl and fentanyl citrate were dissolved in physiological saline for injection. Heroin was provided by NIDA Drug Supply, oxycodone was obtained from Spectrum Chemical MFG Corp (Gardena, CA) and fentanyl was obtained from Cayman Chemical Company (Ann Arbor, MI).

### Surgery

The rats were surgically prepared with chronic indwelling intravenous catheters at ∼13 weeks of age using gas anesthesia and sterile procedures, as previously described (Aarde et al., 2015; Nguyen et al., 2019; Nguyen et al., 2021). Catheters consisted of a 14-cm length polyurethane based tubing (MicroRenathane®, Braintree Scientific, Inc, Braintree MA, USA) fitted to a guide cannula (Plastics one, Roanoke, VA) curved at an angle and encased in dental cement anchored to an ∼3-cm circle of durable mesh. The catheter tubing was passed subcutaneously from a port at the back, inserted into the jugular vein and secured gently with suture thread. A liquid tissue adhesive was used to close the incisions (3M™ Vetbond™ Tissue Adhesive; 1469S B). Two weeks were allowed for surgical recovery prior to starting the experiment. Catheters were flushed with ∼0.2-0.3 ml heparinized (166.7 USP/ml) saline before sessions and ∼0.2-0.3 ml heparinized saline containing cefazolin (100 mg/mL) after sessions. Catheter patency was assessed with the administration of ∼0.2 ml (10 mg/ml) of the ultra-short-acting barbiturate anesthetic, Brevital sodium (1 % methohexital sodium; Eli Lilly, Indianapolis, IN), i.v.. Animals with patent catheters exhibit pronounced loss of muscle tone within ∼3 s of infusion. Animals that failed to display these signs were discontinued from the study and any data that were collected after the previous passing of the test were excluded from analysis. Implantation for one female was not successful, but that animal was retained as a cagemate.

### IVSA Acquisition

Intravenous self-administration (IVSA) was initiated at ∼16 weeks of age in 2 h sessions with 20 µg/kg/infusion heroin available on a Fixed Ratio 1 schedule of reinforcement using procedures previously described (Gutierrez et al., 2025; Nguyen et al., 2019; Nguyen et al., 2018). A response on the drug-associated lever resulted in illumination of a cue light, reinforcer delivery and a 20 second time-out period. A single priming infusion was delivered in a FR session if an animal failed to make a response on the drug-associated lever within 30 minutes of the session start. Sessions were run weekdays (M-F), save for scheduled University holidays. This was continued for 17 total acquisition sessions.

### Dose Substitution

During Sessions 18-22 rats were tested with different per infusion doses of heroin (0, 15, 30, 60, 120 µg/kg/infusion) available on a FR1 schedule of reinforcement in 2 h sessions in a counterbalanced order. This was repeated for doses of oxycodone (0, 30, 60, 150, 300 µg/kg/infusion) on Sessions 23-27 and for fentanyl (0, 0.625, 1.25, 2.5, 5.0 µg/kg/infusion) on Sessions 28-32. Doses were selected based on a prior report which suggested this was sufficient dose range to capture the ascending and descending limbs of a typical inverted U dose-effect function (Wade et al., 2015). One male rat’s catheter failed during the Heroin dose-substitution and thus N=7 males and N=7 females completed these experiments.

Rats were thereafter switched to 3 hour sessions with drug infusions available on the Progressive Ratio Schedule. Doses of heroin (15, 30, 60, 120 µg/kg/infusion; Sessions 33-36), oxycodone (30, 60, 150, 300 µg/kg/infusion; Sessions 37-40) and fentanyl (0.625, 1.25, 2.5, 5.0 µg/kg/infusion; Sessions 41-44) were assessed in a counterbalanced order within drug. One female rat’s catheter failed during the oxycodone PR thus N=7 males and N=6 females completed the oxycodone and fentanyl experiments. Session 45 was used for make-up sessions for three animals due to disconnections of the catheter in prior PR and/or FR sessions.

### Nociception

Latencies for tail withdrawal from 52°C water were assessed after the IVSA dose-substitutions, using methods previously described (Gutierrez et al., 2024; Nguyen et al., 2018). The impact of doses of heroin (0, 0.56, 1.0, 1.56 mg/kg, s.c.) were evaluated with treatment order counter-balanced and 3-4 days between evaluations; a final challenge was conducted with all animals assessed after 2.4 mg/kg, s.c.. This was initiated two weeks after the final IVSA session (mixed doses of fentanyl under a PR schedule) at ∼27 weeks of age. Tail withdrawal latencies were assessed prior to injection and 30 and 60 minutes post-injection.

### Antagonist challenge

All rats with patent catheters were returned to heroin (0.02 mg/kg/infusion) IVSA in two hour sessions under a FR 1 schedule of reinforcement one month after the final PR dose substitution session, ∼30 weeks of age. After six Sessions (45-51), rats were injected five minutes prior to the session with naloxone (0.0, 0.03, 0.3 and 1.0 mg/kg, i.p.), administered in counter-balanced order starting at ∼31 weeks of age. A final challenge was conducted with naloxone 2.0 mg/kg after the completion of the original series.

### Data Analysis

The infusions obtained, percent of responses directed at the drug-associated lever, final ratio completed in the PR procedure and tail-withdrawal latencies were analyzed by 2-way ANOVA including within-subjects factors for acquisition Session, Dose or Pre-Treatment condition, as relevant, and a between-groups factor for Sex. Mixed effects analysis was used in any cases where data points were missing for a given subject. In all analyses, a criterion of P<0.05 was used to infer a significant difference. Significant main effects were followed with post-hoc analysis using Tukey (multi-level factors) or Dunnett (treatments versus control) correction for multiple comparisons. All analysis used Prism for Windows (v. 11.0.0; GraphPad Software, Inc, San Diego CA).

## Results

### IVSA Acquisition

Female rats self-administered more infusions during the first 17 sessions of intravenous self-administration (IVSA) [Sex: n.s., Session: F (4.706, 60.89) = 9.287; P<0.0001, Interaction: F (4.706, 60.89) = 3.690; P<0.01] (**Figure 1**). The Tukey post-hoc test further confirmed significant differences between the sexes for Session 13-17. Priming infusions were delivered to one male in Sessions 4, 8 and 11, and to two males in Sessions 2 and 5. Priming infusions were delivered to one female in Sessions 1 and 4, and to two females in Sessions 7 and 9. The analysis also confirmed a significant effect of Session [F (5.389, 69.39) = 3.26; P<0.01] on the percent of responses on the drug-associated lever and a Dunnett post-hoc test confirmed this discrimination ratio was higher on Sessions 13 and 15 compared with the first session, across groups.

**Figure 1:**
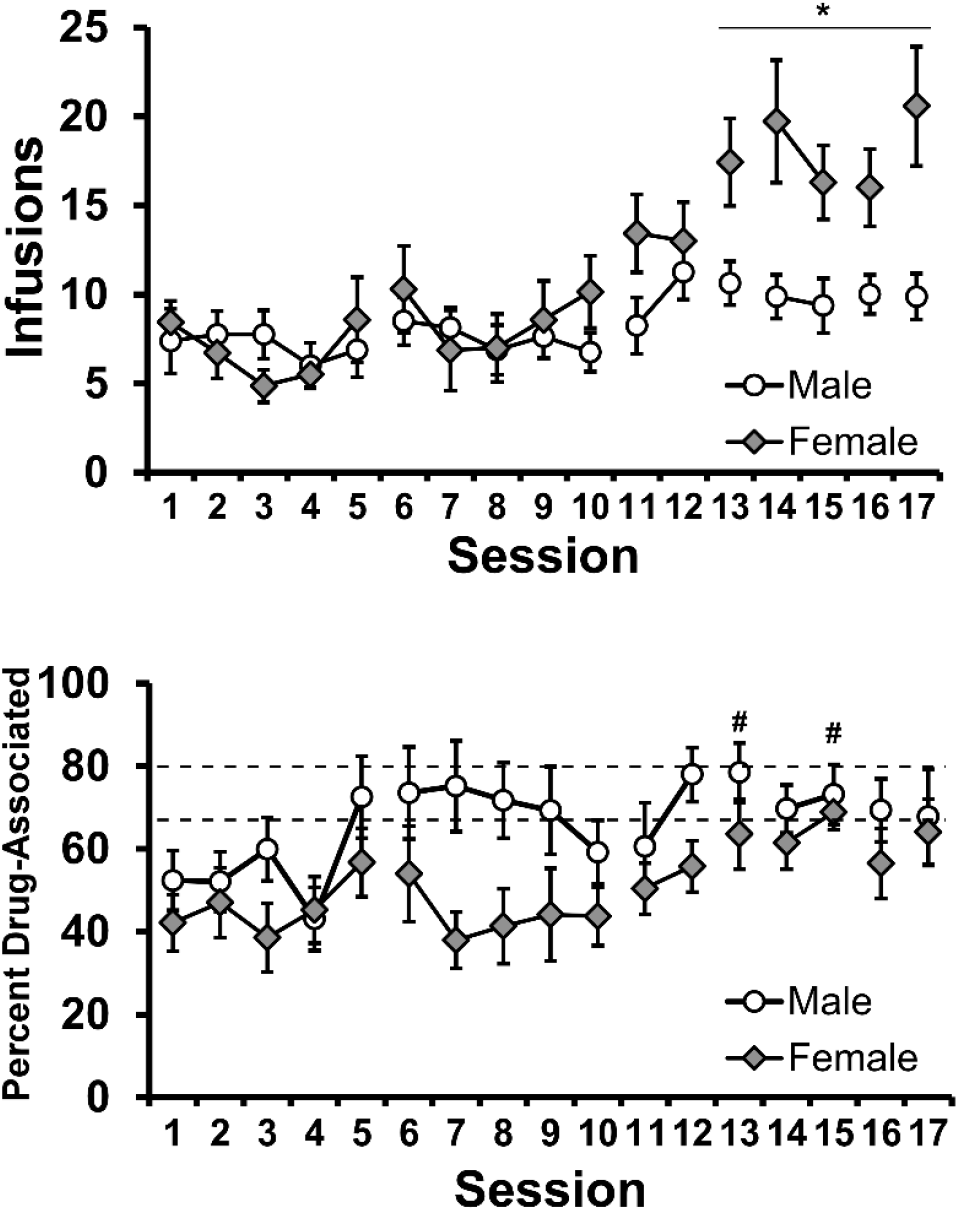
Mean (±SEM) infusions obtained, and percent of responses on the drug-associated lever by female (N=7) and male (N=8) rats for all 17 acquisition sessions. Breaks in the series indicate multiple days between sessions. The dotted lines bracket the 66.7%-80% lever discrimination range for visual comparison. A significant difference between sexes is indicated with * and a significant difference from the first session, across group, is indicated with #.

Examination of mean infusions per 5 minute bin within the session demonstrates a loading dose response in the first five minutes followed by variable rates of responding throughout the session (**Figure 2**). The loading dose ranged from 0-4 infusions in the male rats and 1-5 for the female rats in Session 17 but the group mean was almost identical. There were no more than 2 (males) or 4 (females) infusions obtained by any individual in a given 5 minute bin after the first five minutes.

**Figure 2:**
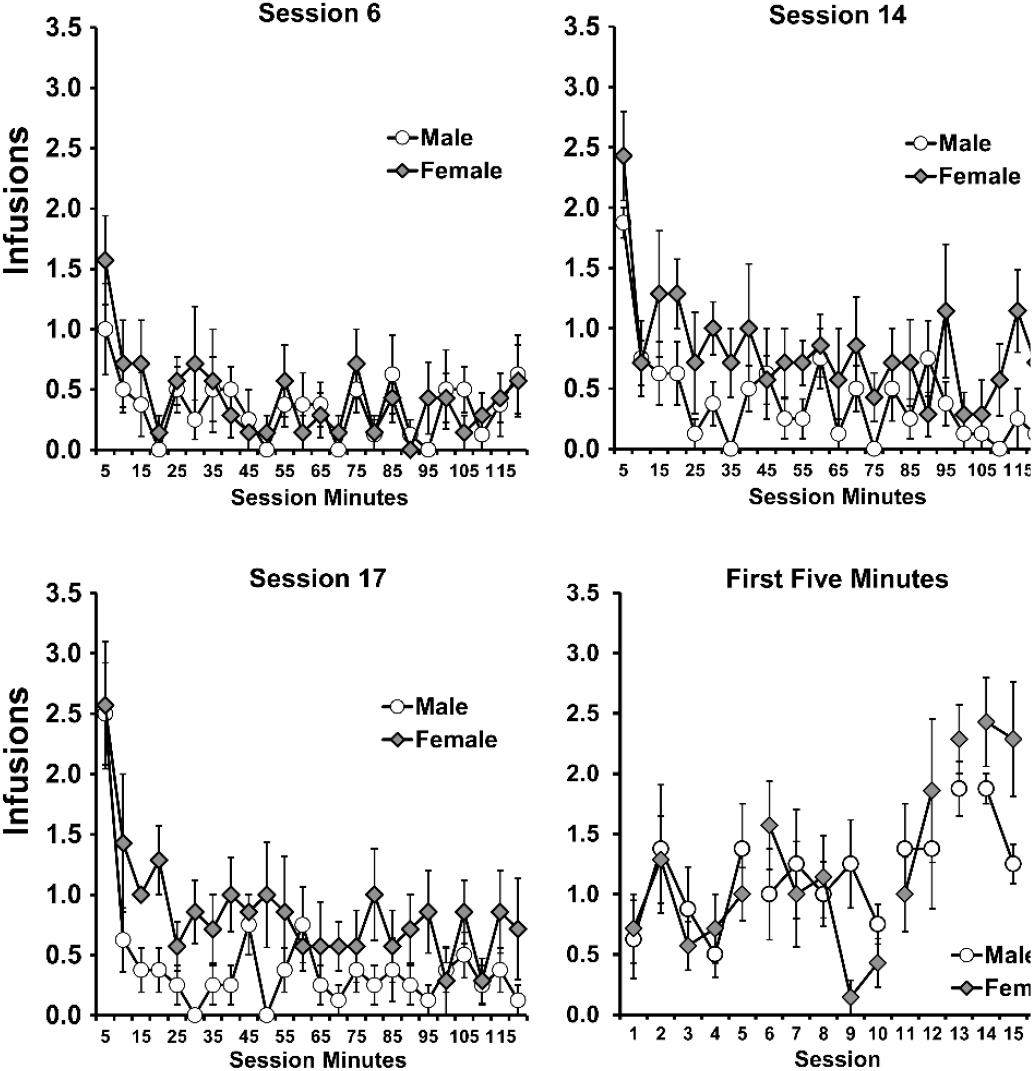
Mean (±SEM) infusions obtained by female (N=7) and male (N=8) rats for each 5 minute interval within the session for Sessions 6, 14 and 17, and in the first five-minute interval for all 17 acquisition sessions.

### IVSA Fixed Ratio Dose-Substitution

There were significant effects confirmed in the analysis of the number of infusions obtained in the heroin [Dose: F (2.459, 29.50) = 12.81; P<0.0001, Sex: F (1, 12) = 7.13; P=0.0204, Interaction: n.s.], oxycodone [Dose: F (2.063, 24.24) = 18.63; P<0.0001, Sex: F (1, 12) = 6.366; P<0.05, Interaction: n.s.] and fentanyl [Dose: F (1.779, 21.35) = 6.11; P<0.01, Sex: F (1, 12) = 4.94; P<0.05, Interaction: n.s.] dose substitution experiments (**Figure 3 A, B, C**). The Tukey post-hoc including all orthogonal comparisons confirmed significant sex differences after individual doses of heroin (60-120 µg/kg/infusion), oxycodone (60-300 µg/kg/infusion), and fentanyl (0, 2.5-5.0 µg/kg/infusion).

**Figure 3:**
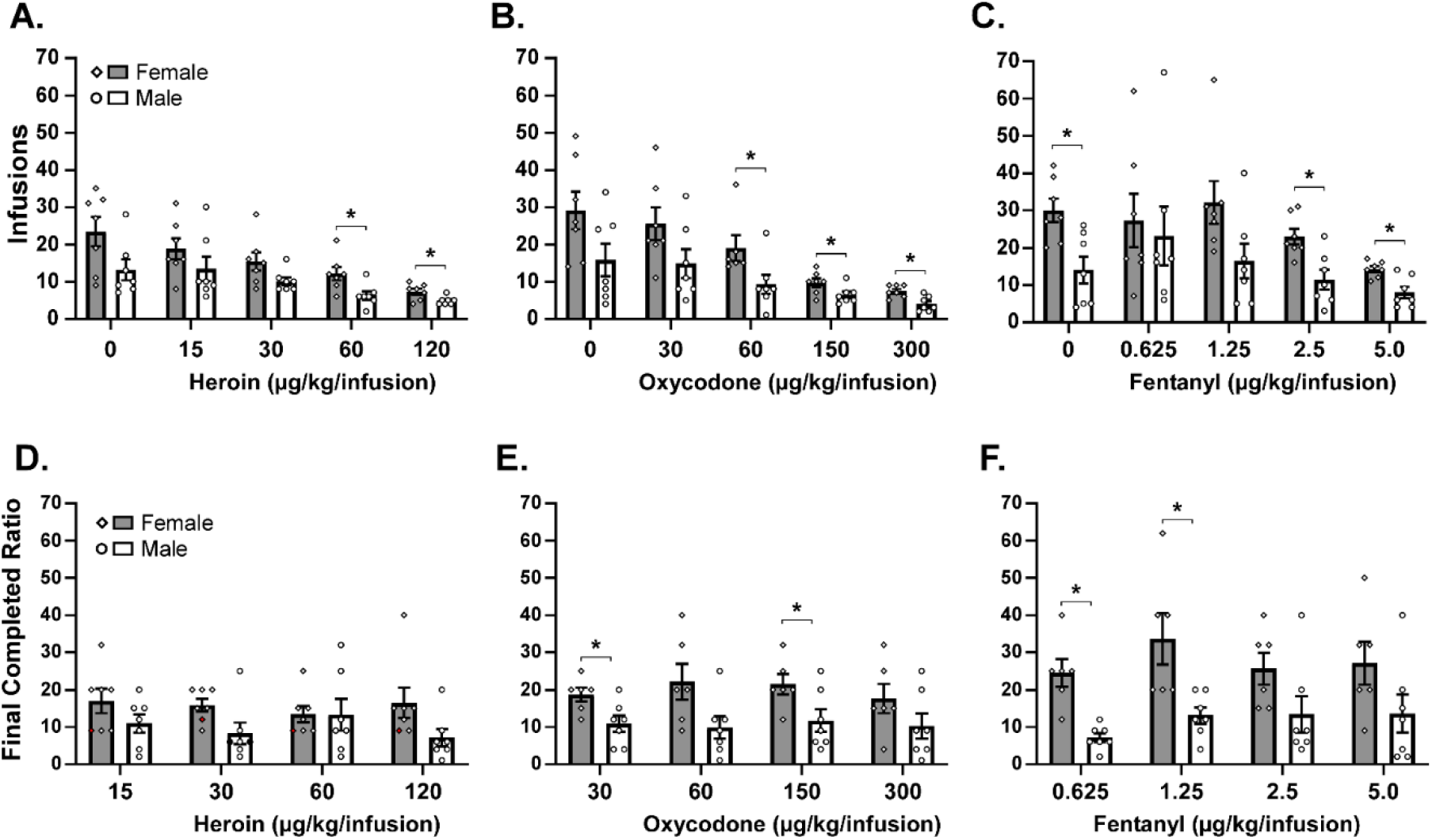
Mean (±SEM) and individual infusions obtained by female (N=7) and male (N=7) rats when A) heroin, B) oxycodone or C) fentanyl doses were available under a FR 1 schedule of reinforcement and mean (±SEM) and individual final ratios completed by female (N=6-7) and male (N=7) rats when D) heroin, E) oxycodone or F) fentanyl doses were available under a Progressive Ratio schedule. A significant difference between sexes is indicated with *.

The Tukey post-hoc analysis of dose, across sex, confirmed significant differences compared with the vehicle condition for heroin (60-120 µg/kg/infusion), oxycodone (150-300 µg/kg/infusion) and fentanyl (5.0 µg/kg/infusion), and compared with the lowest active dose for heroin (60-120 µg/kg/infusion) and oxycodone (150-300 µg/kg/infusion). There were also differences in the self-administration of additional per-infusion doses of heroin (30 µg/kg vs 120 µg/kg), oxycodone (300 µg/kg vs 60 and 150 µg/kg), and fentanyl (5.0 µg/kg vs 1.25 and 2.5 µg/kg) confirmed.

### IVSA Progressive Ratio Dose-Substitution

There were significant effects of sex confirmed in the analysis of breakpoint for oxycodone [F (1, 11) = 6.00; P<0.05] and fentanyl [F (1, 11) = 8.10; P<0.05], but not for heroin (**Figure 3 D, E, F**). The Tukey post-hoc tests further confirmed significant differences between the sexes for oxycodone (30, 150 µg/kg/infusion) and fentanyl (0.625, 2.5 µg/kg/infusion).

### Nociception

The analysis of tail withdrawal latencies confirmed significant effects of Sex (F (1, 14) = 5.10; P<0.05) and Dose (F (2.765, 38.72) = 14.59; P<0.0001), and the Tukey post hoc confirmed significantly longer latencies after heroin (1.0-2.4 mg/kg) compared with vehicle for each sex (**Figure 4**). It also confirmed differences after 1.56-2.4 mg/kg compared the 0.56 mg/kg dose within the female group. The Tukey post-hoc test did not confirm a difference between the sexes at any specific dose.

**Figure 4:**
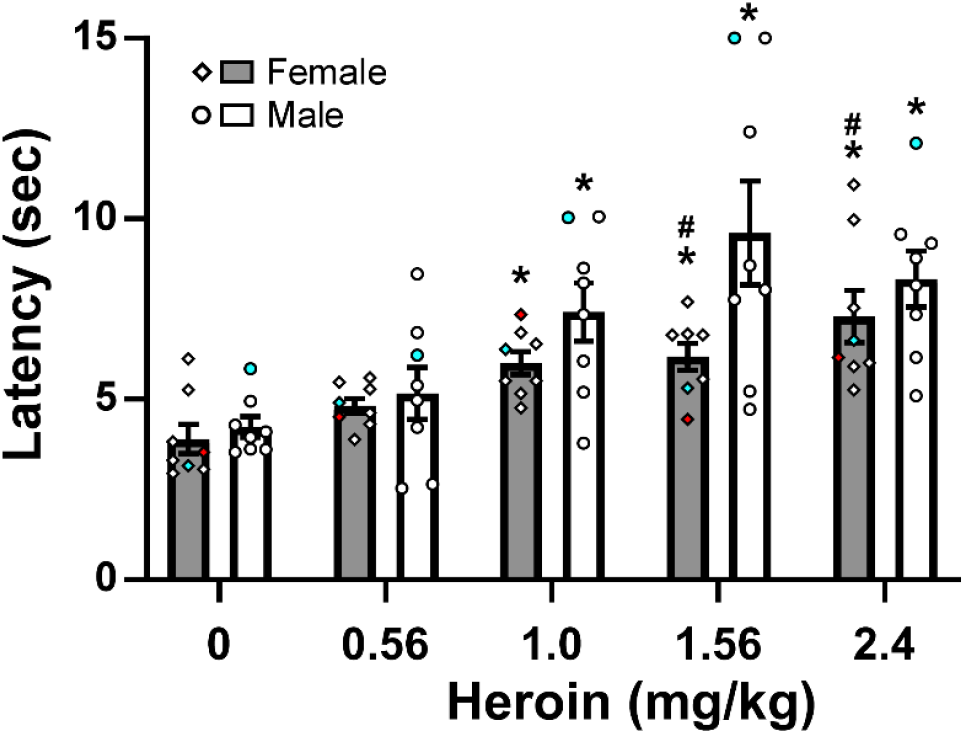
Mean (±SEM) and individual tail withdrawal latencies from female (N=8) and male (N=8) rats following subcutaneous injection with heroin. One female individual did not ever undergo IVSA (red symbol) and one of each sex started IVSA but was discontinued before the others (blue symbols). A significant difference from vehicle is indicated with * and from 0.56 mg/kg with #.

### Antagonist challenge

Administration of naloxone prior to the IVSA session increased the number of infusions obtained (**Figure 5**). The mixed effects analysis confirmed significant effects of Pre-treatment condition [F (2.080, 18.72) = 5.021; P<0.05] and of Sex [F (1, 10) = 18.95; P<0.005]. Post-hoc exploration of the marginal means for pre-treatment condition confirmed a significant difference from the naloxone 1.0 mg/kg condition after 0.03 mg/kg and the 2.0 mg/kg naloxone conditions. The Tukey post-hoc test further confirmed females obtained significantly more infusions in all conditions except 0.3 mg/kg naloxone and within-group the males obtained more infusions after 1.0 mg/kg than they did after 0.03 and 2.0 mg/kg naloxone. One female rat’s catheter was occluded during the baselining sessions, thus N=6 female and N=6 male rats started this experiment; two females did not complete the vehicle condition and one female and one male did not complete the 2.0 mg/kg naloxone condition. Mean lever discrimination of over 80% of responses directed to the drug-associated lever was maintained in both sexes for all treatment conditions, save the males in the vehicle condition (77.1%).

**Figure 5:**
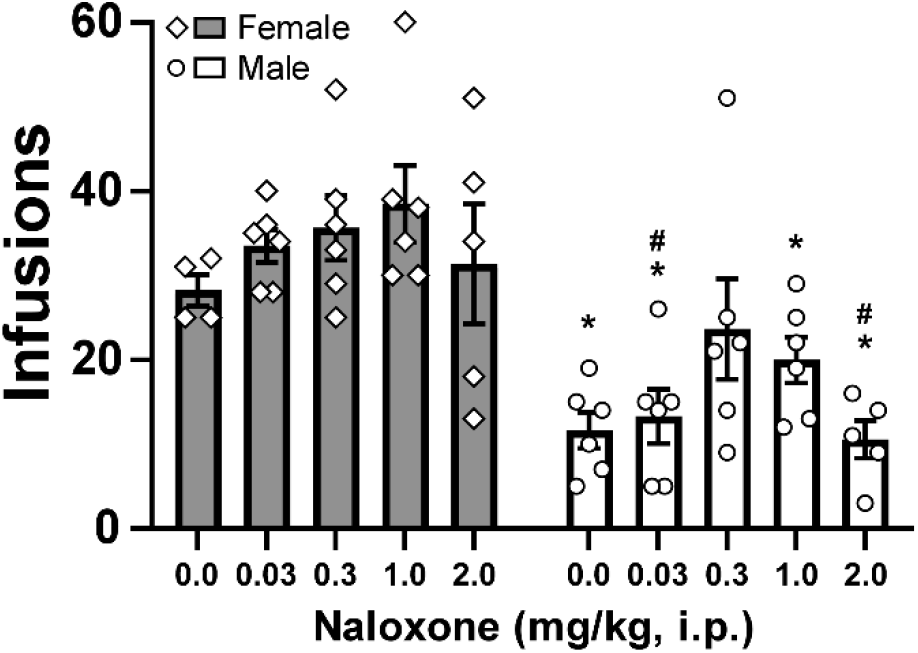
Mean (±SEM) and individual infusions of heroin (20 µg/kg/infusion) obtained by female and male rats under a FR 1 schedule of reinforcement following injection with doses of naloxone (0.0-2.0 mg/kg, i.p.). A significant difference between sexes is indicated with * and a difference within group from the 1.0 dose with #.

## Discussion

The study shows, first, that female heterogenous stock (HS) rats self-administer greater amounts of heroin than do their male counterparts, when operant behavior is reinforced with 20 ug/kg/infusion in 2 h sessions. This sex difference was durable from the end of intravenous self-administration (IVSA) acquisition through additional experimental manipulations of the reinforcing drug (i.e., both dose and opioid identity) under FR and PR response contingencies. That is, female rats self-administered more infusions of heroin, oxycodone and fentanyl in the Fixed-Ratio dose substitution and reached significantly higher breakpoints for oxycodone and fentanyl in the Progressive Ratio test. A blunting of antinociceptive effect of heroin in the female rats was observed after IVSA, suggestive of greater tolerance induced by the greater cumulative self-administered opioid exposure. Female rats’ IVSA was also altered less by pre-treatment with the opioid antagonist naloxone, consistent with drug seeking behavior that is less sensitive to perturbation.

As noted above, there have been a diversity of outcomes in studies which compared the impact of rat sex on opioid IVSA. The present results are consistent with a prior study in which female rats obtained more infusions of morphine in the lower per-infusion doses of a substitution under a Fixed Ratio 4 schedule, and reached higher breakpoints for IVSA of morphine under a Progressive Ratio schedule in 4 h sessions (Cicero et al., 2003). It is also similar to a sex difference observed in Wistar rats trained to self-administer oxycodone in 8 hour sessions and then subjected to dose-substitution (Nguyen et al., 2020). Female rats likewise exhibited greater escalation of heroin intake in a two 3-hr access sessions per day model (Deckers et al., 2025). But the present outcome was dissimilar to a lack of sex-difference in Wistar rats trained to self-administer oxycodone in 6 hour sessions (see (Nguyen et al., 2024); Supplemental Figure S3), and a lack of a sex difference in Sprague-Dawley rats trained in 6 h sessions (Nett and LaLumiere, 2023) or trained first in 2 h sessions followed by two 3 h sessions (D’Ottavio et al., 2023).

Most pertinently, the present results in HS rats are discordant with those of Rakowski and colleagues (Rakowski et al., 2025) who reported no sex differences in the acquisition of IVSA of heroin (20) in ten 2 h sessions by HS rats, and greater intake in the male HS rats when continued post-acquisition on 4 hour daily access. The genetic diversity within the population of HS rats compared with typical laboratory strains cautions that the difference between studies could be the random draw of groups of 8 (present study) or even 28-37 in the Rakowski report. However this study was motivated in part because of a consistent sex-difference observed in the IVSA of heroin (60 µg/kg/infusion) in our lab by Wistar rats and by HS rats selected by genetic prediction of high and low heroin IVSA preference based on a GWAS study (Kuhn et al., 2025a; Kuhn et al., 2025b); not shown. This generalization suggests that training dose and random selection from the genetically diverse HS rat population is unlikely to explain the difference between the present study and the results of Rakowski et al. (2025). There was also no clear evidence that the omission of prior operant training, as with the fluid restriction based pre-training used by Rakowski and colleagues (Rakowski et al., 2025), influenced the outcome. That prior study reported a mean of about 5-6 inactive lever presses by male rats and 4-4.5 by female rats in the final two sessions of acquisition, and roughly 57%-62% of all responses were directed to the drug-associated lever. Similarly, our groups expressed 64% (female) and 68% (male) of responses directed at the drug-associated lever at the end of acquisition. Therefore, the omission of lever training with appetitive reinforcers in our study did not appear to slow acquisition in terms of lever discrimination or the number of reinforcers acquired in the first few acquisition sessions.

Female rats were less sensitive to the anti-nociceptive effects of heroin when assessed after 44 sessions of IVSA. This may be a result of the much higher cumulative exposure throughout the IVSA experiments producing a higher degree of tolerance in the female rats. Alternately, this may be consistent with the IVSA in which females took more infusions on a bodyweight adjusted basis. Although we used a limited range of heroin doses for non-contingent exposure, the 0.56-2.4 mg/kg dose range exceeds the mean self-administered session total in the FR1 highest dose heroin (120 µg/kg/infusion) substitution, i.e., 0.58 mg/kg (males) or 0.88 mg/kg (females), and therefore better reflects (in)sensitivity relative to behavioral preferences. The differences in antinociception between the groups were modest, consistent with a failure to observe consistent sex differences in anti-nociceptive effects of inhaled or injected heroin in our prior work (Gutierrez et al., 2021; Gutierrez et al., 2022; Gutierrez et al., 2020). Interestingly one prior sub-study where sex differences were confirmed found the anti-nociceptive effect was greater in female animals (Gutierrez et al., 2022). A review of the antinociceptive effects of morphine concluded that in most rat strains, included several sources of Sprague-Dawley, males were more sensitive to the anti-nociceptive effects (Mogil et al., 2000). However, a direct comparison of opioid anti-nociception in Sprague-Dawley rats reported an ED_50_ for morphine 2.8 times higher in females, but only a non-significant difference for heroin or oxycodone (0.9 and 1.1 times the male ED_50_, respectively). A similar sex difference for morphine ED_50_, with a reversed sex difference for oxycodone and no sex difference for methadone was also reported (Holtman and Wala, 2006, 2007). This diversity of outcome further emphasizes the potential lability of sex-differences in non-contingent exposure and involuntary responses such as anti-nociception.

In summary, these data show that rat strain, training dose and operant pre-training are not the key differences explaining the lack of sex difference in heroin IVSA acquisition reported in a prior study conducted with HS rats (Rakowski et al., 2025). One potentially explanatory difference is the duration of acquisition since the sex difference in this study only emerged after 10 sessions, i.e., beyond the acquisition interval used in that prior study. Relatedly, it required over 10 sessions for over 80% of male rat groups to meet defined acquisition criterion in prior work (Carroll et al., 2002; Lynch and Carroll, 1999). Overall, this study cautions that the presence or absence of sex differences in opioid IVSA rat models is variable and thus no definitive over-general conclusions should be drawn from any particular model or investigation.

## Acknowledgements

The authors are grateful to Sophia Vandewater, Helen Kim, and Christianne Perral for technical assistance. We are also grateful to Abraham A. Palmer, Ph.D. for provision of HS rats.

